# Functional evolution of olfactory indolergic receptors in Diptera

**DOI:** 10.64898/2026.01.06.697762

**Authors:** Michal Arbel, Esther Yakir, Jonathan D. Bohbot

## Abstract

Indole and skatole are widespread semiochemicals mediating mosquito behaviors such as host seeking and oviposition. These cues are detected by a conserved group of odorant receptors (indolORs) first identified in mosquitoes and later found in brachyceran flies, suggesting that indolic detection may represent a broader dipteran trait. To test this, we identified and functionally characterized ten new candidate receptors from eight species spanning the dipteran phylogeny. Phylogenetic and functional analyses indicate that these receptors form a single, well-supported clade with diversified in sensitivity to indole and skatole. Comparative motif analysis revealed four conserved residues correlated with ligand preference. The previously described indole-sensitive receptor from the lepidoteran *Helicoverpa armigera* OR30 does not exhibit these positions and may have involved sensitivity to indole via another mechanism. These findings underscore the evolutionary trajectory of indolORs across Diptera, revealing both conservation and diversification in indolic odor detection mechanisms.

## Introduction

Insects rely heavily on olfaction to navigate their environment, guiding behaviors such as foraging, mate choice, oviposition, and predator avoidance (Hansson & Stensmyr, 2011). Odorant detection occurs in specialized sensory appendages, most notably the antennae and maxillary palps, which house olfactory sensory neurons (OSNs) within hair-like sensilla (Martin et al., 2011). These neurons express odorant receptors (ORs), which are tetrameric ligand-gated ion channels composed of three conserved obligatory co-receptors (ORco) and a divergent odorant-sensing subunit (ORx) (Wicher et al., 2008). Together, this receptor complex converts environmental volatile organic compounds into electrical signals that the brain translates into behavior. Orco is highly conserved across insect species, typically sharing more than 80% amino acid identity, consistent with its fundamental role in receptor assembly, membrane trafficking, and structural stability (Larsson et al., 2004). In contrast, the odorant-sensing ORx subunits, which mediate ligand recognition, exhibit substantial sequence divergence, with average amino acid identity across species of approximately 20% (Vosshall & Stocker, 2007). Over evolutionary timescales, increasingly complex chemical environments and species-specific ecological pressures have shaped the diversification of insect olfactory systems. For instance, the vinegar fly *Drosophila melanogaster* encodes approximately 62 ORs (Robertson et al., 2003), whereas the yellow fever mosquito *Aedes aegypti* expresses 117 (Matthews et al., 2018) and the common fly *Musca domestica* 86 (Scott et al., 2014), reflecting distinct sensory demands and ecological niches.

Interestingly, despite the extensive diversification of *Orx* genes, a small subset of ORs was identified in mosquitoes, exhibiting unusually high sequence conservation, sharing 50 to 70% amino acid identity (Bohbot et al., 2007a). Functional studies have shown that these receptors respond with high sensitivity and selectivity to indolic compounds such as indole and skatole, and were therefore termed indolORs (Bohbot et al., 2011a; Ruel et al., 2019a). In mosquitoes, indolORs include between two to three paralogs depending on the species: *Or2* and *Or10* are expressed in both Culicinae and Anophelinae (Bohbot et al., 2011a), while *Or9* is restricted to Culicinae and expressed exclusively during the larval stage (Bohbot et al., 2007). Odorant receptor 2 is selectively activated by indole (Bohbot et al., 2011a), whereas OR10 and OR9 are tuned to skatole, with OR9 exhibiting particularly high sensitivity (Ruel et al., 2019a).

Indoles are evolutionarily conserved signaling molecules involved in both intracellular and intercellular communication (J. Lee et al., 2009; J. H. Lee et al., 2015). Synthesized through tryptophan metabolism (Herrmann & Weaver, 1999), they are produced by a wide array of biological sources, including bacteria, fungi, plants, and decaying organic matter. As interspecies and interkingdom cues, indoles mediate diverse ecological interactions among microbes, animals, and plants (Lee et al., 2009). In mosquitoes, indole and its methylated analog, skatole, have been proposed as oviposition attractants, potentially guiding gravid females of species such as *Culex quinquefasciatus*, *Aedes aegypti*, and *Anopheles gambiae* to suitable breeding sites (Beehler et al., 1994; Hughes et al., 2010; Ruel et al., 2019). Beyond oviposition, indole-containing odor blends are also implicated in plant-host attraction, particularly among nectar-feeding mosquitoes like *Ae. Aegypti* (Nyasembe et al., 2014). In non-hematophagous mosquitoes such as *Toxorhynchites amboinensis*, which do not rely on blood meals, indole and skatole detection may facilitate the identification of floral resources or larval food sources (Dekel et al., 2019).

More recently, homologs of mosquito indolORs have been discovered in Brachyceran flies, such as *D. melanogaster* and *M. domestica*, maintaining similar ligand tuning despite lower sequence identity (∼30–43%). Comparative analyses have shown that these brachyceran receptors fall within the same indolOR lineage as the mosquito OR2, OR9 and OR10 families, sharing conserved intron positions, motif architecture and overall phylogenetic placement (Pitts et al., 2021; D. M. Ruel et al., 2021a). These observations raise the possibility that indolORs represent a conserved feature within Diptera. To investigate this further, we functionally characterized indolOR candidates from eight dipteran species spanning both Nematocera and Brachycera, selected to represent a broad phylogenetic and ecological diversity. Using two-electrode voltage clamp (TEVC) electrophysiology in *Xenopus laevis* oocytes, we quantified receptor sensitivity and selectivity toward indole and skatole through EC₅₀ analysis. We also examined *HarmOR30*, a previously identified indole-selective receptor from the moth *Helicoverpa armigera* (Guo et al., 2022), to evaluate whether it shares functional or sequence-level features with dipteran indolORs despite its phylogenetic distance.

## Materials and Methods

### Identification and Phylogenetic Analysis of Dipteran IndolORs

To identify candidate indolOR sequences, we conducted sequence identity searches across the NCBI and VectorBase databases, focusing on twelve dipteran species representing a broad evolutionary spectrum. Species were selected to cover major dipteran lineages and ecological niches, ensuring diverse evolutionary and functional representation. Candidates were selected based on sequence identities ranging from 35% to 47% relative to AaegOR10 **(Fig. 1A).**

**Figure 1.**
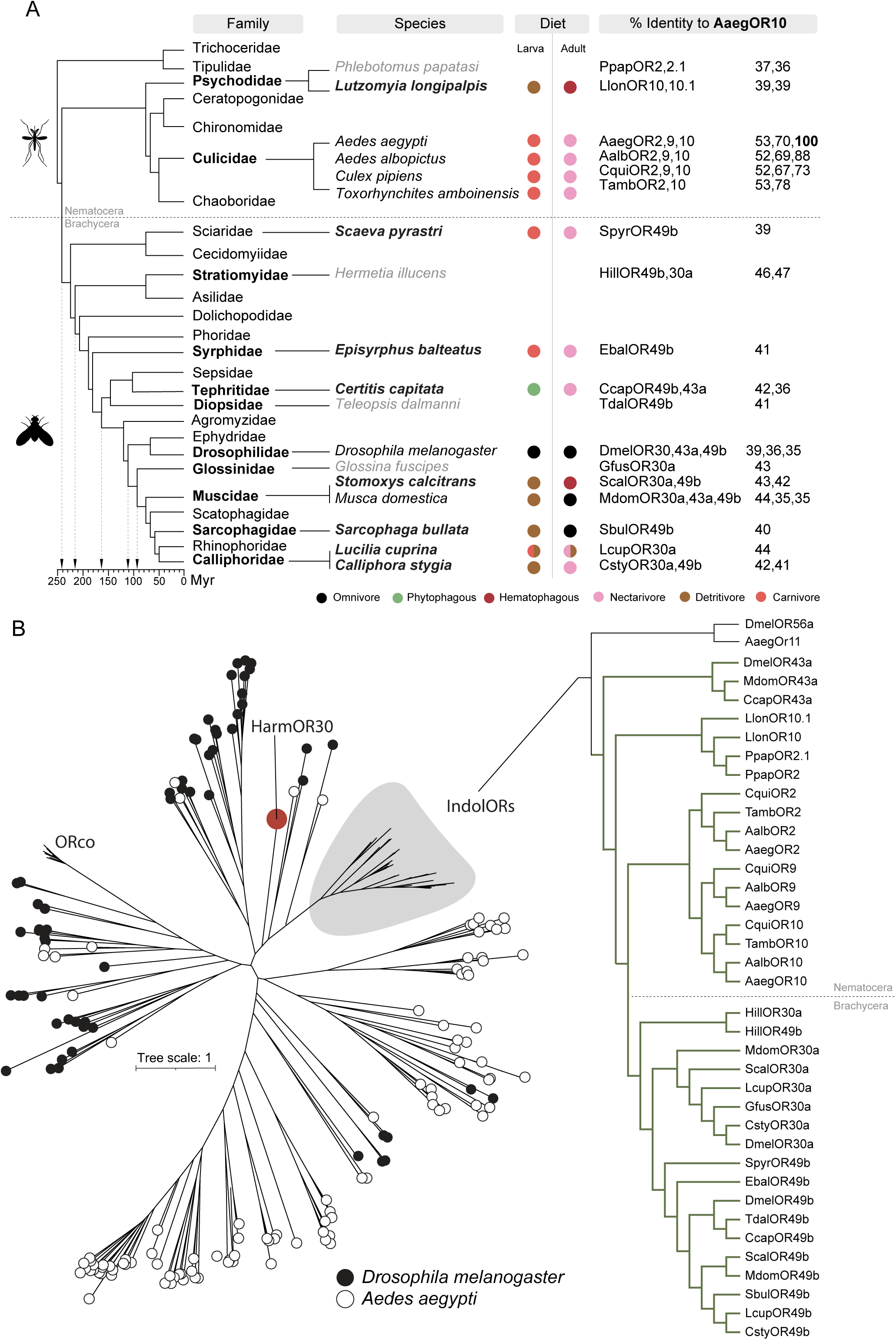
Evolutionary distribution and phylogenetic relationships of dipteran indolORs. (A) Simplified phylogeny of Diptera illustrating the divergence between Nematocera and Brachycera and the diversity of larval and adult feeding ecologies across families, represented by circles denoting larval and adult dietary categories (Adapted from Misof et al., 2014). The species listed represent those in which indolORs have been identified: species newly identified and functionally characterized in this study are shown in bold, whereas species tested here that did not respond to indolic compounds are shown in light grey. The accompanying table summarizes amino acid sequence identity (%) to *Ae. aegypti* OR10. (B) Maximum-likelihood phylogenetic tree of odorant receptors (ORs) from *Ae. aegypti* and *D. melanogaster*, together with all candidate receptors functionally examined in this study. ORs from *Ae. aegypti* are highlighted in white, *D. melanogaster* ORs in black, and *Helicoverpa armigera* OR30 in red. The tree shows that all newly identified receptors cluster within the dipteran indolOR clade, forming distinct nematoceran and brachyceran sublineages (green), whereas *HarmOR30* segregates outside this group, consistent with its divergent tuning and motif composition.

For the phylogenetic analysis, amino acid sequences of all candidate indolORs identified in this study were used, together with previously characterized indolORs from the literature and the full OR repertoires of *Ae. aegypti* and *D. melanogaster*. This dataset enabled the construction of a comparative framework to evaluate the evolutionary relationships among indolORs and the broader OR family.

Sequences were aligned using MAFFT version 7 (Katoh and Standley 2013). A maximum-likelihood phylogenetic tree was constructed with IQ-TREE software (Bazinet et al., 2014), using the best-fit model selected automatically by the software. Bootstrap analysis with 1,000 replicates was performed to assess the robustness of the resulting tree **(Fig. 1B, Supplementary Fig. 1)**. The tree was visualized and edited using the Interactive Tree of Life (iTOL) software. An unrooted layout was selected to avoid implying a specific evolutionary direction, but the tree was oriented by positioning the conserved ORco sequences as a visual reference. Although ORco was used to help organize the tree for interpretability, the tree remains mathematically unrooted. The sequences analyzed included 106 ORs from *Ae. aegypti* (including the indolORs AaegOR2, AaegOR9, and AaegOR10), 64 ORs from *D. melanogaster* (including DmelOR30a, DmelOR43a, and DmelOR49b), and all candidate indolORs from the dipteran species listed in this study (**Fig. 1A**).

### IndolOR Gene Cloning

Candidate genes encoding indolORs (**Supplementary Table 1**) were codon-optimized and synthesized by Twist Biosciences. The cloning followed previously established methods (Dekel et al., 2019; Ruel et al., 2021a). Briefly, synthesized genes were inserted into the pENTR entry vector using the Gateway directional cloning system and subcloned into the pSP64t-RFA expression vector for in vitro transcription. Plasmids were purified using the GeneJET Plasmid Miniprep Kit, and the integrity of the coding sequences was confirmed by Sanger sequencing. Complementary RNAs (cRNAs) were synthesized from linearized pSP64t-RFA vectors using the mMESSAGE mMACHINE SP6 kit.

### Odorant Preparation

Indole and skatole were dissolved in 200 µL of DMSO to produce a 1 M stock solution, which was further diluted in ND96 saline buffer supplemented with 0.8 mM CaCl2. These compounds were chosen due to their demonstrated potency as ligands in previous studies on indolergic receptors. Both indole and skatole have been identified as highly effective in activating indolORs across various species, with skatole showing particularly strong binding affinities in species like *Ae. aegypti* and *D. melanogaster* (Ruel et al., 2019; Ruel et al., 2021). Odorant concentrations used to establish concentration-response relationships were formulated through dilution of the initial 1 M stock.

### Two-Electrode Voltage Clamp of Xenopus Oocytes

To characterize the ligand responsiveness and tuning profiles of candidate dipteran indolORs, we employed the TEVC technique using *Xenopus laevis* oocytes. This method allows precise measurement of receptor-mediated ion currents in response to odorant stimulation, making it highly suited for comparing the functional sensitivity and selectivity of indolORs across species. Stage V–VI oocytes were harvested, enzymatically treated with 1 mg/mL collagenase at 18°C for 2 hours at 70 RPM, then washed five times in Ca²⁺-free Ringer solution, rinsed in a washing solution, and washed five additional times with incubation medium supplemented with 5% dialyzed horse serum, 50 μg/mL tetracycline, 100 μg/mL streptomycin, and 550 μg/mL sodium pyruvate. Oocytes were allowed to recover overnight before microinjection.

To facilitate efficient functional characterization across multiple species, we employed a cross-species OR–ORco pairing strategy. This approach leveraged the high degree of evolutionary conservation of the ORco subunit across insects and was supported by previous studies demonstrating functional compatibility between ORs and heterologous ORcos (Larsson et al., 2004; Zhao et al., 2024). We selected *Hermetia illucens* ORco (HillORco) for most receptor pairings, with *Ceratitis capitata* and *Lutzomyia longipalpis* as the only exceptions where their native ORco was used. Validation experiments pairing *Aedes aegypti* OR9 (AaegOR9) with HillORco yielded an EC50 of 12.6 nM **(Supplementary Fig. 2)**, closely aligning with the 5 nM reported for the native AaegORco (Ruel et al., 2019a), thereby confirming the functional reliability of this method. This strategy allowed us to efficiently evaluate receptor function across multiple species without compromising experimental accuracy.

Oocytes were injected with 27.6 nL of RNA (receptor + co-receptor) using a Nanoliter 2010 injector (World Precision Instruments, Inc., Sarasota, FL, USA) and incubated at 18°C for three days. Recordings were conducted using an OC-725C oocyte clamp (Warner Instruments, LLC, Hamden, CT, USA) at a holding potential of −80 mV. Oocytes were placed in an RC-3Z recording chamber and exposed to 8-second pulses of odorant solution in increasing concentrations of indole or skatole. Whole-cell currents were recorded using a Digidata 1550A interface with pCLAMP10 software (Molecular Devices, Sunnyvale, CA, USA). Between stimulations, oocytes were washed to allow currents to return to baseline. The current amplitude measured at each concentration point was systematically recorded to derive EC₅₀ values, providing precise quantitative measures of receptor sensitivity.

### EC₅₀ Values and Statistical Treatment

All statistical analyses were performed using GraphPad Prism 8 Software (GraphPad Software Inc., La Jolla, CA, USA). Dose-response data were analyzed using nonlinear regression with a four-parameter logistic model to derive EC50 values. The Hill slope was constrained to 1 to fit a standard dose-response curve. The goodness of fit was evaluated using R² values, ensuring that the derived EC50 values reliably represented the receptor-ligand interactions. Significance of differences between indole and skatole responses was evaluated by F-tests of model fits. For previously published receptors, EC₅₀ values were extracted from the literature; in several cases statistical comparisons were provided (typically t-tests), while in others only relative EC₅₀ differences could be inferred. These receptors were nevertheless included, as all showed clear and substantial separation in sensitivity between the two ligands **(Supplementary Table 2).**

### Motif and Structural Analysis

Motif discovery was conducted using MEME v5.5.8 (Bailey et al., 2015) in discriminative mode to identify short conserved sequence motifs enriched in dipteran indolORs relative to other dipteran ORs. The input set comprised 25 dipteran indolOR protein sequences, and the background set consisted of 650 non-indolORs representing the complete OR repertoires of the same species with indolORs excluded. MEME was configured to search for up to 15 motifs with widths of 6 to 8 amino acids, using a zoops model (zero or one occurrence per sequence) and a zero-order Markov background model (**Supplementary Fig. 3**).

Motif enrichment was subsequently evaluated using SEA (Simple Enrichment analysis) with a significance threshold of p < 1 × 10⁻⁴ (**Supplementary Table 3**). The frequency of motif occurrence in indolORs and background ORs was tested using Fisher’s exact test, and *p* values were adjusted using the Benjamini–Hochberg procedure to control the false discovery rate. For downstream analyses, we focused on the nine motifs detected in all 25 dipteran indolORs, corresponding to MEME-2 to MEME-10, which represent the most consistently conserved features of the indolOR subfamily.

To assess how these indolic motifs are distributed in receptors outside the indolOR group, we scanned three outgroup ORs using FIMO (Find Individual Motif Occurences) with the indolOR-derived position weight matrices. AaegOR11 and DmelOR56a were included because they cluster near the dipteran indolOR clade in the phylogeny, although they do not exhibit canonical indole or skatole tuning. HarmOR30 from *H. armigera* was included because it responds to indole but falls outside Diptera. Motif matches with *p* < 1 × 10⁻⁴ were recorded together with their positions. This approach allowed all three outgroup receptors to be evaluated using a fixed motif set that was defined solely from confirmed indolORs (**Supplementary Table 3**).

For each receptor, M2–10 motifs were expanded into their constituent residues, generating a complete and gap-free alignment of equivalent positions across all indolORs (**Supplementary Table 4**). Each position was then evaluated for its contribution to tuning differences between indole- and skatole-sensitive receptors. To assess the quantitative impact of each position on ligand selectivity, we compared the difference in ligand sensitivity (ΔEC₅₀,) across residue states. Analyses were limited to receptors with available EC₅₀ data (n = 22). Positions containing two residue states were tested using Mann–Whitney U tests, and those with three or more states using Kruskal–Wallis tests. Corresponding non-parametric effect sizes were calculated as Cliff’s δ (two states) or ε² (multi-state), quantifying the proportion of variance in ligand preference explained by each site. Multiple-testing correction was again applied using FDR (**Supplementary Table 5**).

Categorical associations between residue identity and receptor tuning were tested using contingency tables: Fisher’s exact tests were applied for 2×2 comparisons and χ² tests for larger tables (**Supplementary Table 5**). *p* values were corrected for multiple testing using the Benjamini–Hochberg false discovery rate (FDR). All statistical analyses and visualizations were performed in Python 3.11, employing the pandas, scipy.stats, and seaborn libraries.

## Results

### Phylogenetic Analysis Reveals a Conserved IndolOR Clade Across Dipteran Species

To place the newly identified receptors (**Supplementary Table 1**) in an evolutionary context, we performed a maximum-likelihood phylogenetic analysis based on their amino acid sequences. The tree included all candidate receptors from our search, previously characterized indolORs, and the full OR repertoires of *Ae. aegypti* and *D. melanogaster*. We also included HarmOR30, a lepidopteran receptor previously reported as narrowly tuned to indole and expressed in the proboscis of *H. armigera* (Guo et al., 2022), to explore whether indole-responsive receptors exist outside Diptera and whether they share evolutionary links with dipteran indolORs. The resulting topology revealed a well-supported indolOR cluster (bootstrap 99.9), clearly separated from other OR lineages. Within this cluster, we recovered the major sublineages previously described (Bohbot et al., 2011; Pitts et al., 2021; Ruel et al., 2021): the Nematocera-specific OR2, OR9, and OR10 clades (bootstrap 97.4) and the Brachycera-specific OR30a and OR49b groups (bootstrap 99.8). The OR43a lineage formed a distinct basal sublineage within the indolOR cluster (bootstrap = 100), branching prior to the divergence of Nematocera and Brachycera, supporting its designation as an ancient member of the dipteran indolOR family. HarmOR30 also clustered within the broader indolOR clade, supported by a bootstrap value of 78.1, but did not fall within any of the established dipteran sublineages (**Fig. 1B and supplementary Fig. 1**).

### Dipteran IndolORs Exhibit Lineage-Specific and Conserved Tuning Profiles

Building on the phylogenetic analysis, we functionally characterized candidate indolORs to evaluate their responses to indole and skatole. We screened candidate indolOR from 12 dipteran species (**Fig. 1A**) from which ten ORs responded to indole and/or skatole (**Fig. 2**). Using TEVC recordings in *X. laevis* oocytes (**Fig. 2A**), we measured odorant-evoked current responses across increasing concentrations, generated concentration–response curves (CRCs), and derived EC₅₀ values, providing a detailed and quantitative assessment of receptor sensitivity towards indole and skatole (**Fig. 2B**). For all receptors characterized in this study, EC₅₀ values for indole and skatole were significantly different as determined by F-tests, with the exception of CstyOR49b, which gave a *p* value of 0.0524 (**Supplementary Table 2**).

**Figure 2.**
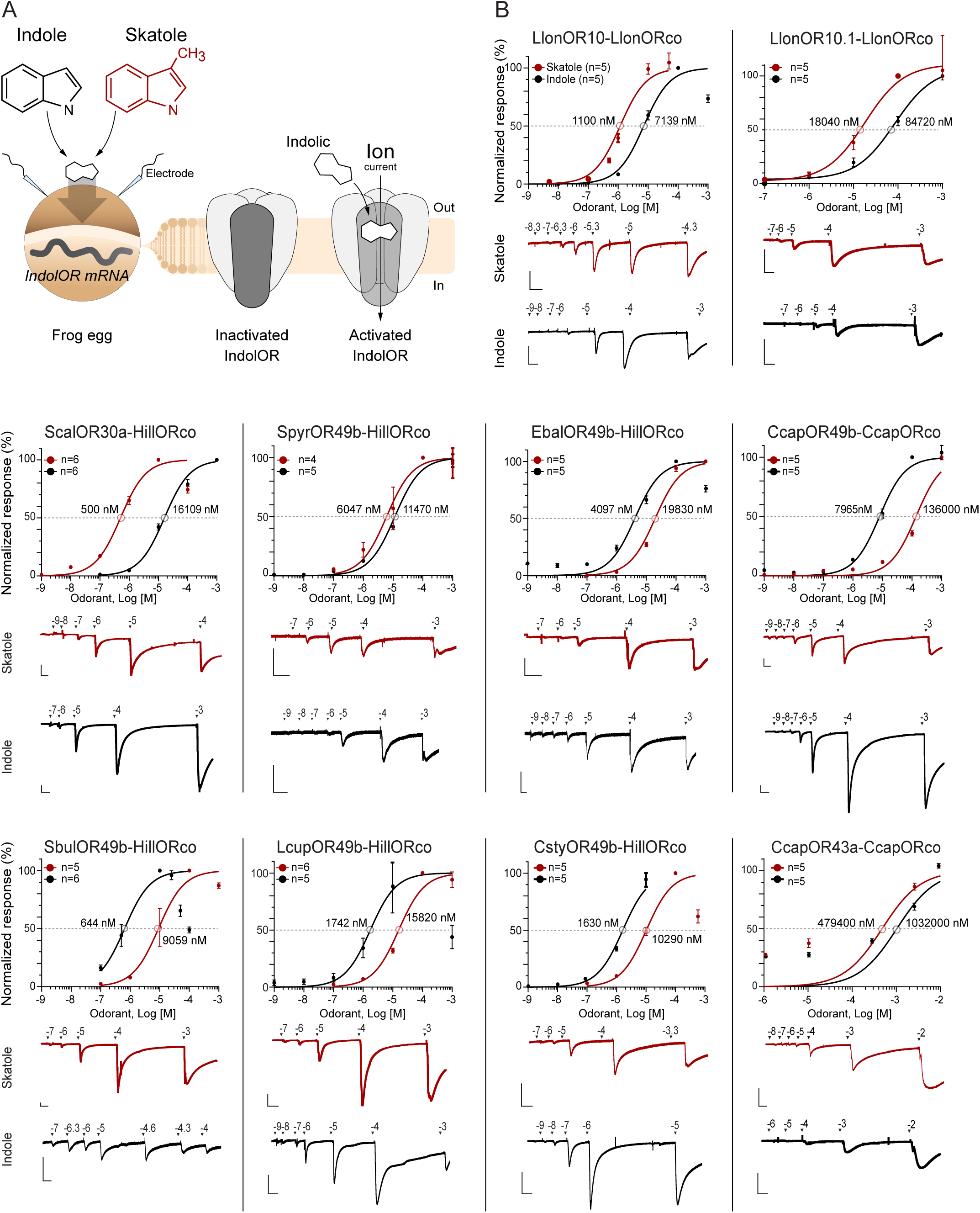
Functional characterization of dipteran indolORs in *Xenopus laevis* oocytes. (A) Schematic illustration of two-electrode voltage-clamp (TEVC) recordings used to measure indolOR responses. mRNA encoding an indolOR is injected into *X. laevis* oocytes, where the receptor is expressed in the membrane. Exposure to indolic ligands, indole or skatole, induces receptor activation and opening of the ion channel, producing measurable inward currents. Indole and skatole share a common indole ring backbone, differing only by a methyl substitution, which underlies their closely related but distinct receptor activation profiles. (B) Normalized dose–response curves for ten candidate indolORs co-expressed with either their native ORco or *Hermetia illucens* ORco (HillORco). Interpolated EC₅₀ values for skatole (empty red dots) and indole (empty black dots) are indicated on each curve. Data points represent mean ± SEM (n = 4–6 oocytes). Representative current traces are shown below each panel; scale bars represent 200 nA and 1 minute.

We observed a range of receptor sensitivities to indole and skatole, with some ORs matching previously characterized indolORs and others exhibiting distinct tuning profiles **(Fig. 3)**. Among the Nematocera, the only species from which we successfully expressed and recorded receptors was the New World sand fly *Lutzomyia longipalpis* (**Fig. 2B**, **Fig. 3**). The two identified homologs (LlonOR10 and LlonOR10.1) responded more strongly to skatole than indole. Overall, LlonOR10 was more sensitive than LlonOR10.1 towards both ligands.

**Figure 3.**
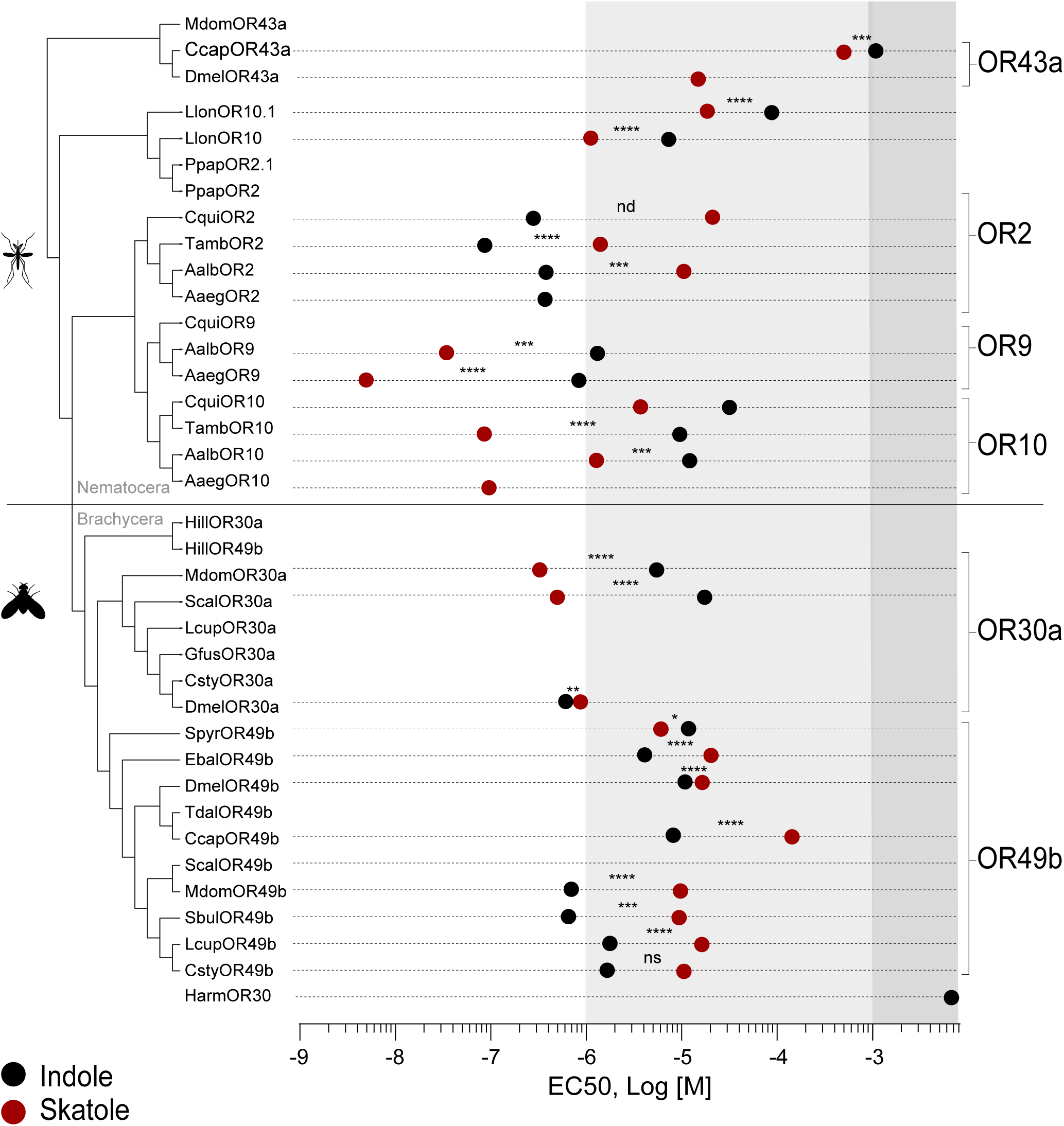
Sensitivity and tuning profiles of dipteran indolORs. Half-maximal effective concentrations (EC₅₀) of newly characterized and previously reported indolORs. Solid black dots represent responses to indole and solid red dots represent responses to skatole. Each receptor is arranged phylogenetically according to its corresponding indolOR lineage (OR2, OR9, OR10, OR30a, OR49b, OR43a). Asterisks denote statistical significance between indole and skatole sensitivities based on pairwise comparisons: *p* < 0.05 (*), *<* 0.01 (**), < 0.001 (***), and < 0.0001 (****); ns = not significant, nd = not determined.

In Brachycera, a larger set of receptors was functionally characterized. *Stomoxys calcitrans* OR30a showed a tuning profile similar to *Musca domestica* OR30a, with higher sensitivity to skatole in the nanomolar range and lower sensitivity to indole in the low-to-mid micromolar range. Within the OR49b group, tuning profiles varied. Receptors such as *Scaeva pyrastri* OR49b and *Episyrphus balteatus* OR49b displayed mid-to-high micromolar EC₅₀ values for both indole and skatole, similar to *D. melanogaster* OR49b. In contrast, OR49b homologs from *Calliphora stygia*, *Lucilia cuprina*, and *Sarcophaga bullata* exhibited higher sensitivity to indole (nanomolar to low micromolar EC₅₀ values), and weaker responses to skatole, aligning with the tuning profile of *Musca domestica* OR49b. A single new representative of the OR43a group was characterized from *Ceratitis capitata*. This receptor exhibited high EC₅₀ values for both indole and skatole, indicating relatively low sensitivity to either ligand. HarmOR30 responded to indole (EC₅₀ of 228.9 µM) but not to skatole consistent with a previous report (Guo et al., 2022) (**Fig. 4**).

**Figure 4.**
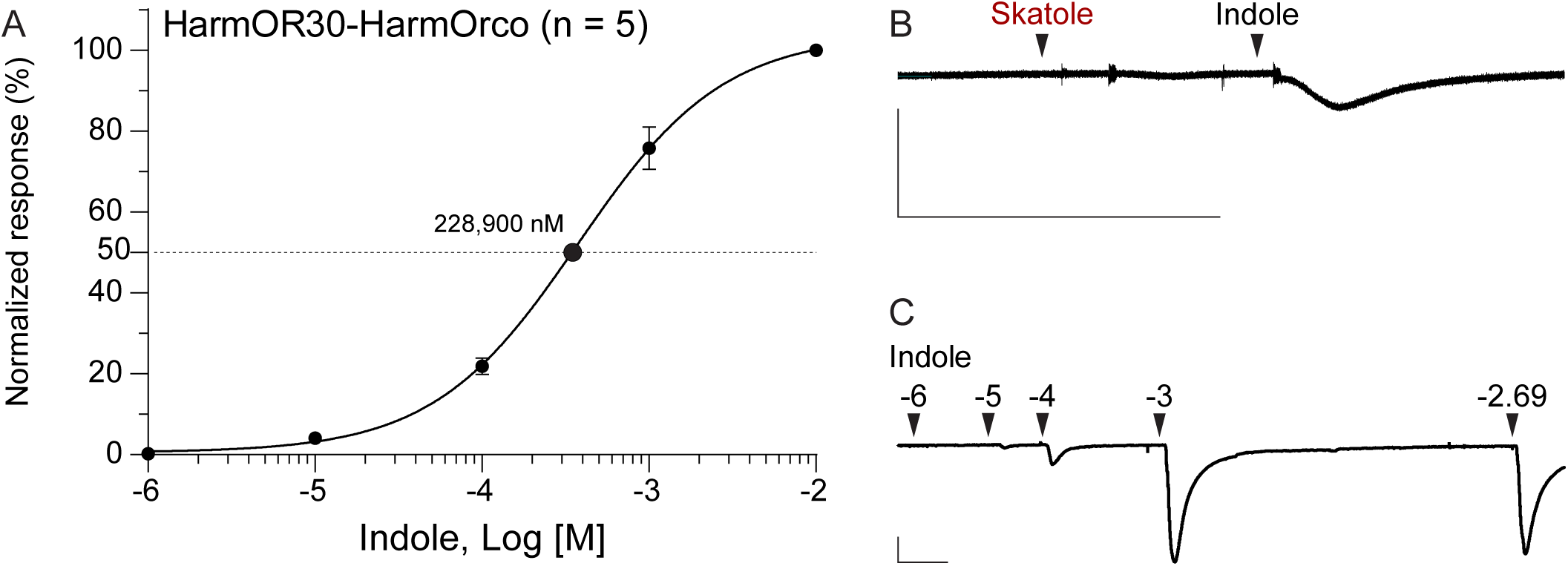
Functional characterization of *Helicoverpa armigera* OR30 (HarmOR30). (A) Normalized concentration–response curve for indole (n = 5; EC₅₀ ≈ 229 µM). (B) Representative trace showing responses to 100 µM skatole and indole. (C) Representative traces to increasing indole concentrations. Scale bars: 200 nA, 1 min.

### Motifs Associated with Dipteran IndolORs Ligand Selectivity

We hypothesized that the conserved sequence motifs shared across dipteran indolORs contribute to selectivity between indole and skatole. We first identified conserved short amino-acid sequence elements across all indolORs and then examined whether they correlated with indolic selectivity. Motif discovery using MEME revealed 15 conserved motifs among the 25 indolOR sequences (Supplementary Fig. 3). Using SEA to assess enrichment against a background of 650 non-indolORs confirmed that all 15 motifs were significantly enriched (Supplementary Table 3). Of these, nine motifs (M2–M10) appeared in all receptors. Motif 1, which had the strongest statistical support of all motifs (E-value 5.7 × 10⁻¹²⁴), was detected in every dipteran indolOR but was absent from the OR43a-family of receptors and was therefore excluded from subsequent analyses. Structural mapping of motifs M2-M10 onto the AaegOR10 cryo-EM model (Zhao et al., 2024) **(Fig. 5A)** showed that motifs M3, M5 and M9 corresponded to regions involved with ligand interactions. Motif 5 and 9 lie along helices S6 and S2, respectively, forming opposing walls of the binding pocket, whereas M3 is located on the extracellular loop connecting S3 and S4 near the pocket entrance. The remaining six motifs were positioned in regions distant from the predicted binding pocket, such as helices S0 and S5, and the C-terminal domain that forms part of the ion channel gate and is enriched in interaction sites with Orco. Although these motifs are less likely to interact directly with ligands, they were still included in the subsequent analyses to allow a comprehensive assessment of conserved sequence features across the indolORs.

**Figure 5.**
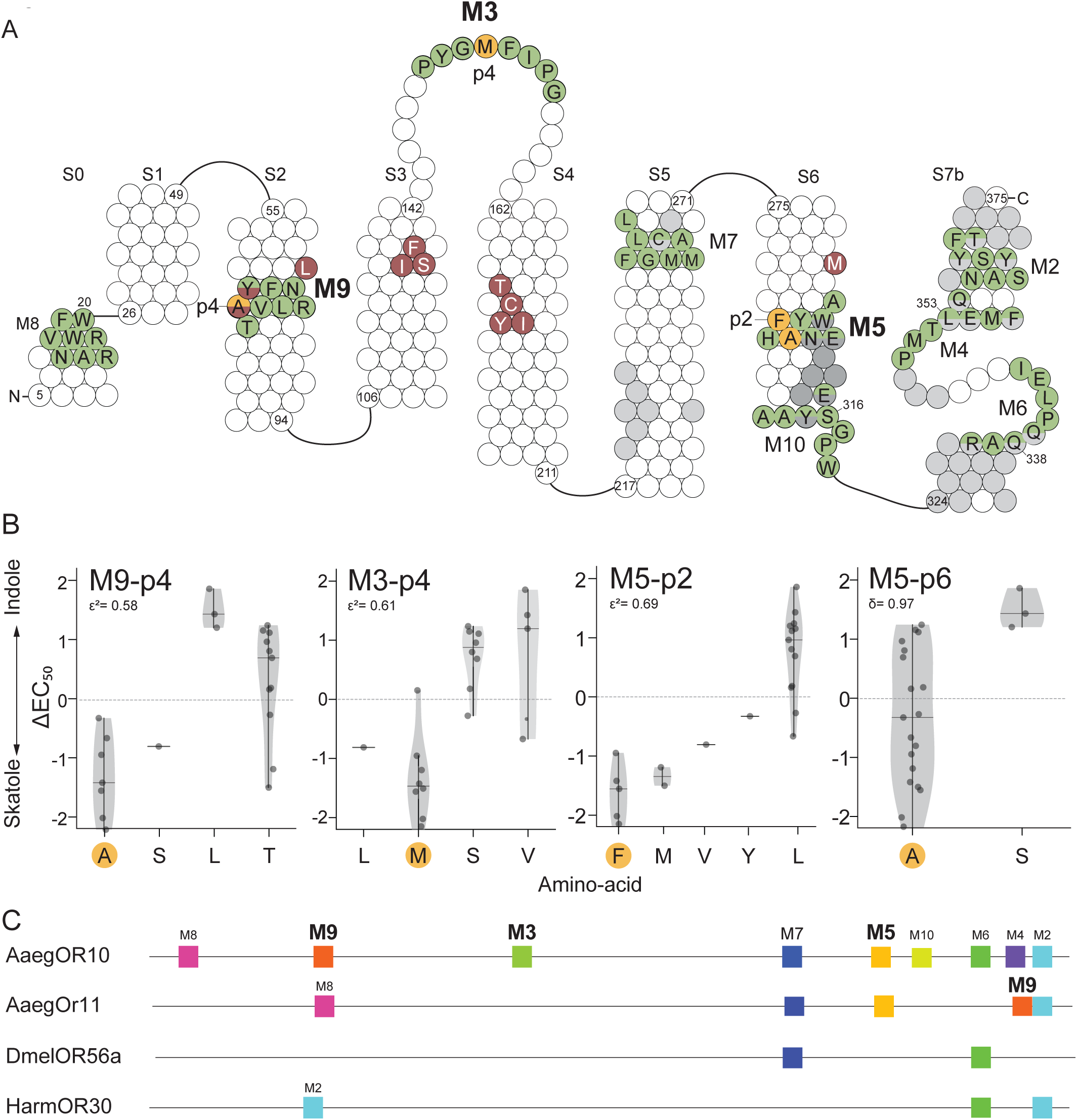
Conserved indolic motifs and fine-tuning positions in dipteran indolORs. (A) Topology map of AaegOR10 based on the *o*-cresol–bound cryo-EM structure reported by Zhao et al. (2024). All nine conserved indolOR motifs are shown in green. Positions with demonstrated functional impact on *o*-cresol activation, including the three validated pocket residues L67, S133 and Y183, are indicated in bown. Residue L67 contributes a hydrophobic pocket contact, S133 hydrogen-bonds to the *o*-cresol hydroxyl, and Y183 provides an aromatic pocket interaction, with alanine substitutions at each site reducing o-cresol affinity. Residues reported in the structure to form contacts with ORco are shown in grey. The four positions with the strongest quantitative effects on ligand selectivity (M3-p4, M5-p2, M5-p6 and M9-p4) are shown in yellow. (B) Violin plots showing the relationship between residue identity at the four fine-tuning positions and ΔEC₅₀. Effect sizes (ε² or δ) are indicated for each position, and highlighted residues correspond to AaegOR10. (C) Comparison of conserved indolic motifs across an indolOR representative (AaegOR10), two dipteran ORs that cluster closely with indolORs (AaegOR11 and DmelOR56a), and a non-dipteran candidate indolOR (HarmOR30).

We next asked whether specific residues within these motifs correlate with ligand selectivity across receptors with known EC₅₀ values for both indole and skatole. For each aligned position within motifs M2-M10 (**Supplementary Table 4**), we tested the relationship between residue identity and ΔEC₅₀ = log EC₅₀ (skatole) – log EC₅₀ (indole) using non-parametric rank tests (Mann–Whitney U for two-state, Kruskal–Wallis for multi-state positions), with effect sizes expressed as Cliff’s δ or ε² and FDR correction for multiple testing (**Supplementary Table 5**). Four positions exhibited significant associations with ΔEC₅₀, all located in or near the binding pocket (**Fig. 5B**). Position 4 in motif 3 (M3-p4) lies on the S3–S4 loop near the cavity entrance, where methionine was linked to greater skatole sensitivity, whereas serine and valine were associated with indole sensitivity. In motif 5, located within the transmembrane helix S6 forming one wall of the pocket, two positions (M5-p2 and M5-p6) emerged as potentially influential. At M5-p2, leucine correlated with higher indole selectivity, while phenylalanine and methionine were linked to skatole selectivity. At M5-p6, alanine and serine showed a similar pattern, being associated with skatole and indole selectivity, respectively. M9-p4, on helix S2 forming the opposite pocket wall, showed that leucine was mainly associated with a high preference for indole, alanine with a strong preference for skatole, while threonine appeared in both receptor types. Three of these positions (M3-p4, M5-p2, and M9-p4) also showed significant categorical enrichment between tuning classes (**Supplementary Table 5**).

To evaluate how the indolic motif architecture extends to receptors outside the indolOR clade, we examined AaegOR11 and DmelOR56a, the closest neighboring receptors identified in our phylogeny **(Fig. 1B)**. AaegOR11 is most strongly activated by the terpenoid cis-decahydro-1-naphthol, with weaker responses to several indolic compounds (Pullmann-Lindsley et al., 2024), whereas DmelOR56a is narrowly tuned to geosmin (Yao, 2025). Scanning AaegOR11 using FIMO identified five of the nine conserved indolic motifs (M2, M5, M7, M8 and M9) **(Fig. 5C**, **Supplementary Table 6**). Three motifs were aligned with their corresponding positions in dipteran indolORs, while two others were found in different locations. Motif 9 occurred at positions 400 to 407 within S7b rather than at the conserved S6 pocket-adjacent region characteristic of indolORs, and M8 appeared in the region that corresponds to the indolOR M9 site but with weak statistical support. Among the three motifs associated with the binding-pocket environment, only M5 was present and correctly positioned, while M3 was not detected. In DmelOR56a, FIMO identified only two conserved motifs, M6 and M7 **(Fig. 5C**, **Supplementary Table 6**). Both were located in regions associated with Orco interactions and match their positions with dipteran indolORs. None of the pocket-associated motifs M3, M5 or M9 were present **(Fig 5C)**. We also scanned HarmOR30, in which only motifs M2 and M6 were detected. Motif 2 was identified twice (positions 66–73 and 369–376), and M6 at positions 344–351. Both M2₃₆₉–₃₇₆ and M6 are found within the conserved C-terminal region that interacts with Orco, whereas M2₆₆–₇₃ occurs far from the corresponding M2 position in dipteran indolORs and likely represents a spurious match **(Fig 5C**, **Supplementary Table 6**).

## Discussion

Our findings support the hypothesis that indolORs represent a conserved feature within Diptera. By functionally testing 19 receptors from 12 species using two-electrode voltage clamp assays, we identified 10 previously uncharacterized indolORs from 8 species spanning both Nematocera and Brachycera. These assays revealed a consistent lineage-associated tuning profile, with most receptors responding to indole or skatole in the low micromolar range.

In Nematocera, *Lutzomyia longipalpis*, our sole newly tested representative, encoded two candidate indolORs that cluster within the indolOR clade of the phylogeny but do not group with any previously defined mosquito sublineage. LlonOR10 displayed a low micromolar response consistent with the nematoceran indolOR profile, whereas LlonOR10.1 exhibited a higher EC_50_ values than other nematoceran skatole receptors, suggesting that it may be preferentially activated by other, more potent indolic ligands that remain to be identified.

In Brachycera, most indolORs we tested exhibited EC₅₀ values between approximately 0.5 and 10 micromolar for either indole or skatole, matching the sensitivity range previously described in *Drosophila melanogaster* and *Musca domestica*. The only exception was CcapOR43a, which showed markedly lower sensitivity, consistent with the tuning characteristics reported for DmelOR43a. As noted in previous studies (Pitts et al. 2021; Ruel et al. 2021), the OR43a lineage branches at the base of the indolOR clade, before the split between Nematocera and Brachycera, indicating that this group represents an early-diverging lineage that retains elements of ancestral indolic responsiveness.

These observations suggest that dipteran indolORs are defined not only by a conserved phylogenetic identity and intron architecture but also by their low-sensitivity responses to indolic compounds and by a complete indolergic motif triad consisting of M3, M5 and M9. This triad forms the structural foundation of the dipteran indolOR architecture, while coordinated substitutions at a small number of positions embedded within it refine ligand preference between indole and skatole. The lineage-specific response patterns we observed raise the question of how these functional differences are encoded at the structural level, particularly given the clear phylogenetic coherence of the indolOR family. To address this, we examined the conserved motif architecture that defines dipteran indolORs and evaluated how variation within this framework correlates with ligand selectivity. Across our dataset of indolOR motifs, coordinated substitution patterns at these four positions are associated with ligand-selectivity. A central feature of dipteran indolORs is the conserved arrangement of nine short motifs distributed across the receptor. Three of these, M3, M5 and M9, occupy fixed positions around the predicted ligand-binding cavity and form a core structural triad in indolORs. These motifs lie on the S3–S4 extracellular loop (M3) and the S6 and S2 transmembrane helices (M5 and M9), which surround and shape the binding pocket. To determine whether specific residues within this structural framework influence tuning to indole or skatole, we mapped EC₅₀ values from our TEVC dataset onto conserved motif positions. The tuning analysis identified four positions associated with indolic selectivity, all located within the three motifs that surround the binding pocket. Positions M5-p2 and M5-p6 are positioned inside the binding cavity on the S6 helix. M9-p4 lies on the S2 helix and was shown in the AaegOR10 cryo-EM structure to interact with o-cresol, providing direct structural evidence for its role in ligand placement. Moreover, in *Cx. quinquefasciatus* OR10, mutations at this position reverse ligand preference between indole and skatole (Franco et al., 2022). M3-p4 lies on the S3–S4 loop at the pocket opening, raising the question of whether loops are involved in ligand selectivity. For example, modifications in loop residues can markedly alter odorant potency, as demonstrated for the human receptor OR1A1 (Geithe et al., 2017).

The ORs phylogenetically nearest to the indolOR clade further illustrate this structural pattern. AaegOR11, which is activated only by high concentrations of indolics, retains M5 in the correct location, while M3 and M9 are absent or not aligned with their canonical positions. DmelOR56a, a geosmin receptor with no detectable indolic sensitivity, lacks all three indolergic motifs (M3, M5 and M9). HarmOR30, despite its specificity for indole, lacks the full motif triad and exhibits substantially reduced responsiveness compared to dipteran indolORs. Taken together with its phylogenetic separation from Diptera, these features suggest that HarmOR30 has independently acquired indolic sensitivity through a distinct molecular mechanism.

This study establishes a comparative structural and functional framework for understanding indolic detection in Diptera. Several limitations should be acknowledged. Only two indolic ligands were tested, and several receptors did not exhibit measurable responses, which may be attributable to limited expression or the absence of the appropriate activating ligands among those tested. Expanding the chemical panel and sampling additional dipteran families will help refine the extent and variability of the indolOR scaffold. Sequence-based identification of indolORs proved reliable within Diptera but did not predict indolic sensitivity outside Diptera, including Hymenoptera, Coleoptera, Orthoptera, Mecoptera and Siphonaptera, the closest sister groups to Diptera. These groups represent promising targets for future work aimed at identifying whether indolORs are present outside Diptera or whether the indolOR scaffold represents a dipteran-specific adaptation.

## Supplementary Figures

**Supplementary Figure 1. Phylogenetic placement of candidate indolORs.** Maximum-likelihood phylogenetic tree of odorant receptors (ORs) from *Ae. aegypti*, *D. melanogaster*, and all candidate receptors functionally investigated in this study. Branch support values are indicated by bootstrap scores (1,000 replicates). The shaded grey region marks the dipteran indolOR clade, showing strong statistical support for the grouping of newly identified receptors within this lineage. The HarmOR30 is a homologous (red area) but clearly more distantly related indolOR.

**Supplementary Figure 2. Validation of *Hermetia illucens* ORco (HillORco) as a functional co-receptor for cross-species assays.** (A) Normalized dose–response curve of *Aedes aegypti* OR9 coexpressed with HillORco in *Xenopus laevis* oocytes (n = 5). The EC₅₀ for skatole was 12.8 nM, closely matching the 5 nM value reported for AaegOR9 with its native ORco (Ruel *et al*., 2019). (B) Representative current traces showing responses of AaegOR9-HillORco to increasing concentrations of skatole. Scale bars: 200 nA, 1 min. These results confirm the functional compatibility of HillORco and validate its use as a universal co-receptor for heterologous indolOR-ORco pairings in this study.

**Supplementary Figure 3. MEME motif discovery analysis of dipteran indolORs.** Output of the MEME Suite motif discovery analysis showing the locations and consensus sequences of 15 motifs (motif-1–motif-15) identified across 25 dipteran indolOR sequences. Colored blocks indicate motif positions within each receptor. The p value shown for each receptor corresponds to the MAST sequence-level significance score, reflecting the overall match of the receptor sequence to the inferred motif set. Structural region annotations above the diagram indicate extracellular loop and transmembrane segment occupancy of motifs, based on the cryo-EM structure of AaegOR10. These motifs were used for subsequent comparative and quantitative analyses to identify conserved regions associated with indolic compound detection and fine-tuning.

## Supplementary Tables

**Supplementary Table 1.** List of candidate dipteran indolORs.

**Supplementary Table 2.** EC₅₀ values and statistical comparison of indole and skatole responses.

**Supplementary Table 3.** SEA (Simple Enrichment Analysis) for conserved indolOR motifs.

**Supplementary Table 4.** Motif sequence matrix for indolORs.

**Supplementary Table 5.** Position-level statistical analysis of motif residues.

**Supplementary Table 6.** FIMO motif occurrence analysis in outgroup receptors (AaegOR11, DmelOR56a) and the lepidopteran HarmOR30

## Acknowledgments

This study was supported by ISF 719/21 awarded to J.B.

## Author contributions

MA conducted all experiments. MA and JB conceived the study. EY provided genetic constructs and injectable mRNA. MA and JB wrote the manuscript.

