## Supplementary Figures for "Functional evolution of olfactory indolergic receptors in Diptera"

### Supplementary Figure 1

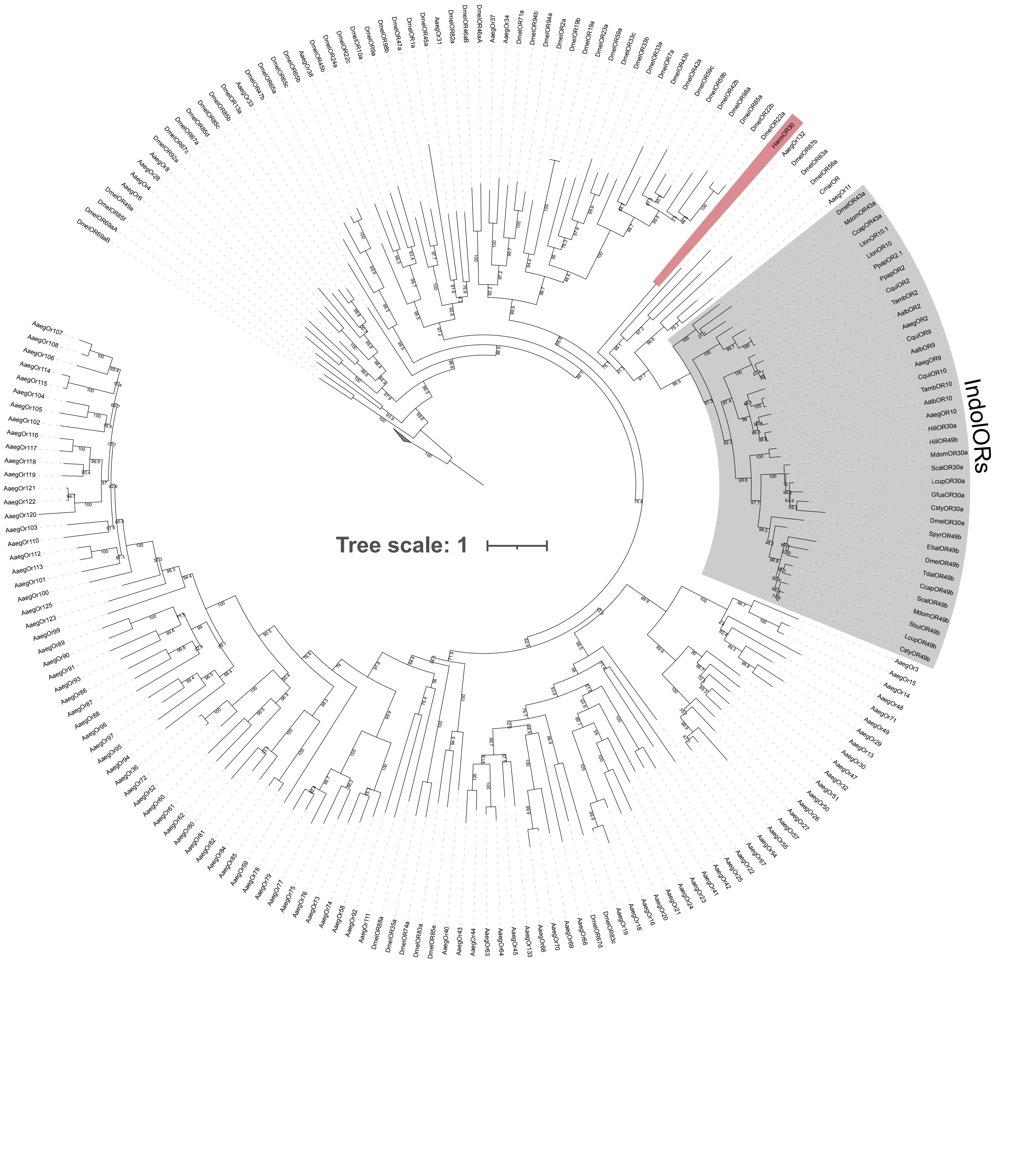

Supplementary Figure 2

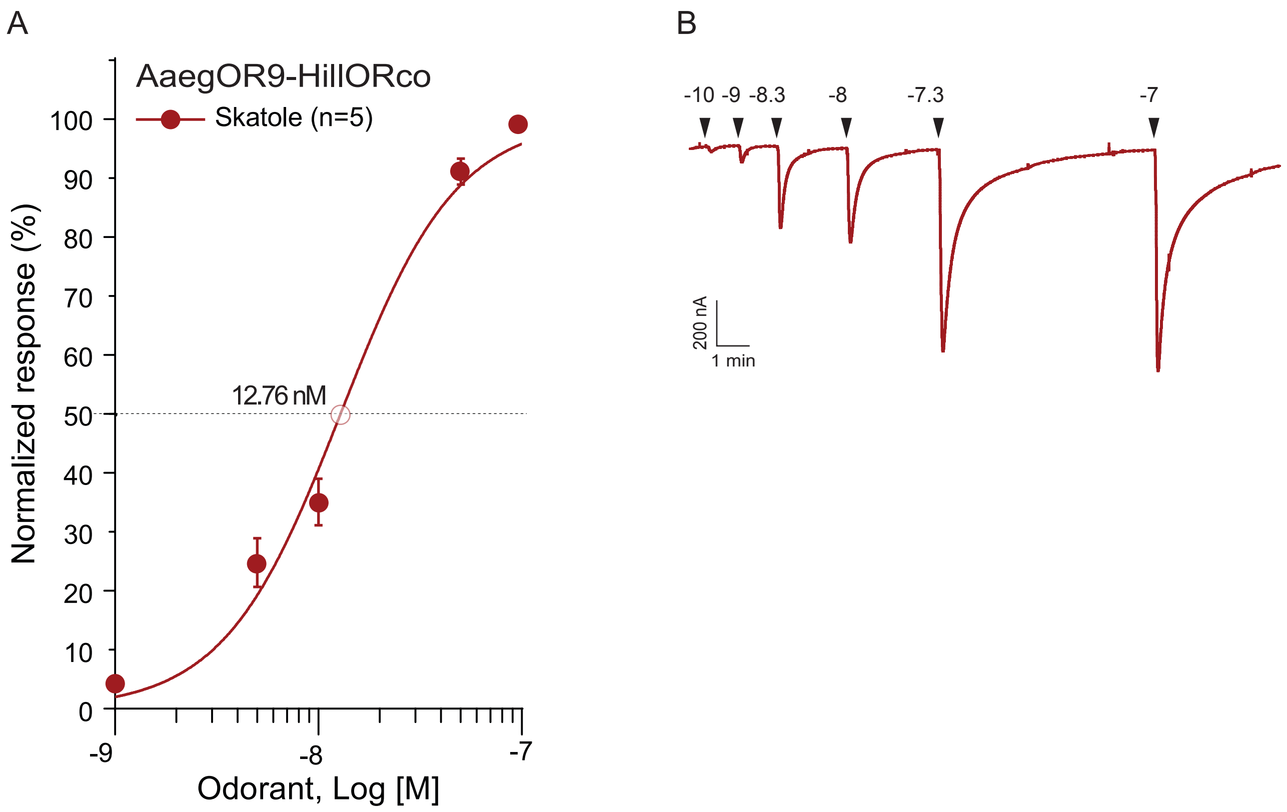

#### Supplementary Figure 3

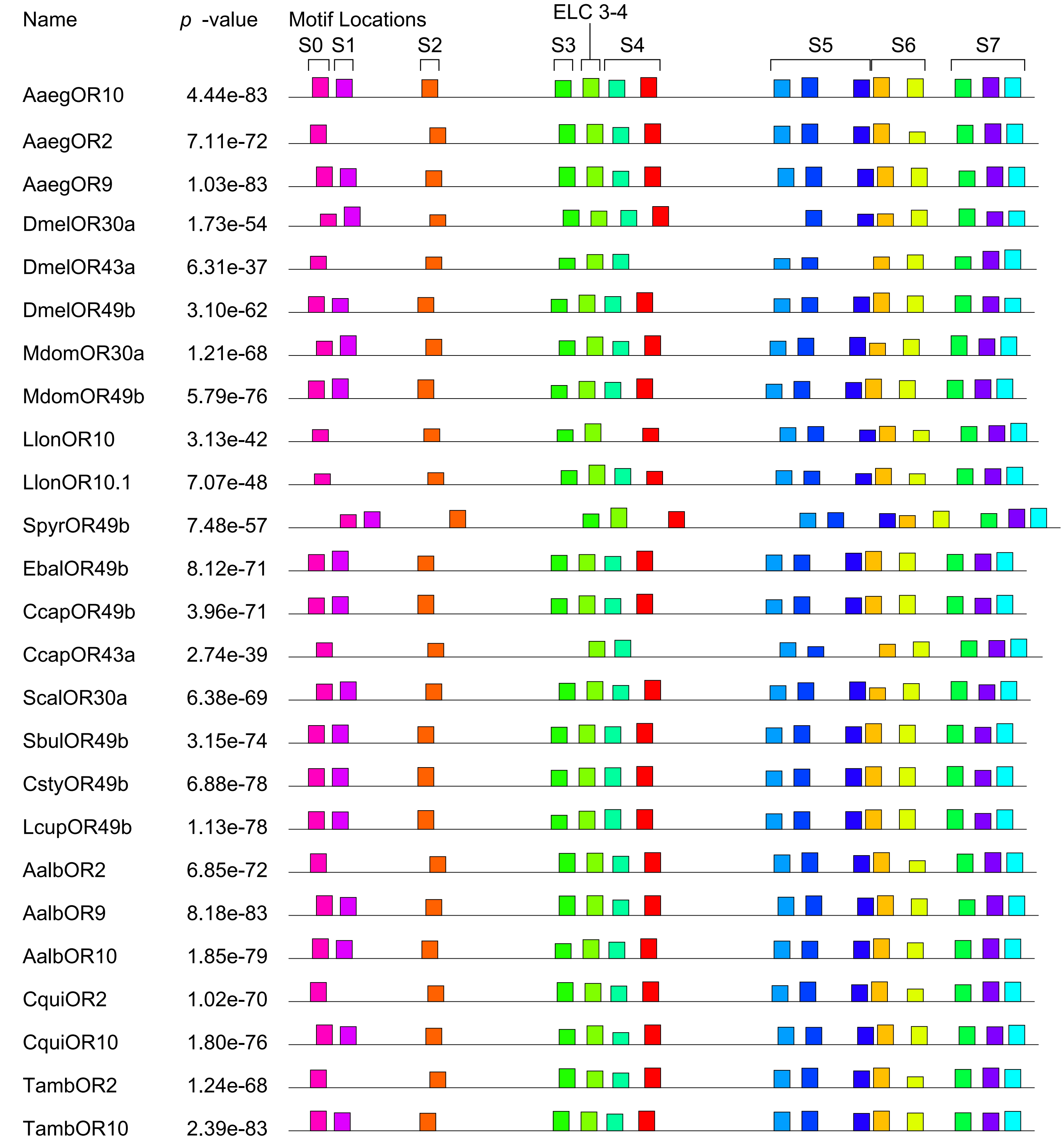


| Motif Symbol | Motif Consensus |
|---|---|
| 1 | PGCCMYIP |
| 2 | NASYSYFT |
| 3 | PYGMYIPG |
| 4 | PMTLEMFQ |
| 5 | ALYWHANE |
| 6 | RAQKPLEI |
| 7 | FGMMLCAL |
| 8 | NVRVWKFW |
| 9 | YFNTVLRA |
| 10 | EAAYETPW |
| 11 | GFVVYPLF |
| 12 | SPFYEIWY |
| 13 | NELVTYIF |
| 14 | YIFMIFSQ |
| 15 | DHBWRRYI |
